# Mammal extinctions and the increasing isolation of humans on the tree of life

**DOI:** 10.1101/154526

**Authors:** Sandrine Pavoine, Michael B. Bonsall, T. Jonathan Davies, Shelly Masi

**Author notes:** **Author for correspondence:** Sandrine Pavoine,.

## Abstract

A sixth great mass extinction is ongoing due to the direct and indirect effects of human pressures. However, not all lineages are impacted equally. As humans, we frequently believe that we hold a unique place on Earth. Here, we show that our current impacts on the natural world risk to heighten that expectation. Evolutionary proximity to *Homo sapiens* emerges as a powerful predictor of extinction risk among mammals. Our analysis shows that the species most closely related to *H. sapiens* are exposed to a large variety of threat types and that they may also have greater intrinsic sensitivity to threats. Pruning back the tree of life around us will lead to our species being among those with the fewest close relatives. We will erase our evolutionary history, forcing its uniqueness. If no action is taken, we will lose crucial biodiversity for the preservation of Earth ecosystems, and a key reference to what makes us human.

## 1. Introduction

Earth is experiencing a major extinction crisis [1]. Marine and terrestrial ecosystems are undergoing profound changes as a result of human activities. Indeed, if primary causes for past mass extinction events still remain unclear, the anthropogenic cause of the current biodiversity crisis is widely understood [2]. The loss of biodiversity is usually evaluated in terms of number of species driven extinct. In particular, about one-fifth of vertebrate species are threatened with extinction [3] and 75% of current vertebrate species could be lost in the following centuries [1]. Yet, species are not equivalent, and not all lineages are impacted equally.

Phylogenetic diversity provides an alternative measure of biodiversity which captures the shared evolutionary histories of a set of species. Yet, phylogenetic diversity is often poorly protected by current conservation practices which target species-rich areas with high endemism, but ignore species relatedness [4]. Losing even a single species may represent the loss of millions of years of unique evolutionary history. However, future projections are even more dramatic: as extinction risks are clustered in the tree of life, entire lineages could disappear [4-6], with consequences for the stability of ecosystems [7].

Human impacts vary markedly across taxa. The extinction risks of mammals have been particularly well studied compared to other groups [8]. Mammals include more than 5,000 species. They are charismatic and perform important and essential ecosystem functions. About a quarter of mammals are threatened with extinction [8]. Some mammalian groups of species, however, are more threatened (e.g. primates, Perissodactyla, Cetartiodactyla) than others (e.g. Chiroptera and Rodentia) [5].This phylogenetic clumping in extinction risk is predicted to lead to higher loss of evolutionary history than previously thought, at least in these groups [4, 5].

Among the most endangered lineages, it is particularly striking that more than half of primate species, our closest relatives, are currently under threat [9]. Here, we analyze an underexplored potential correlate of species extinction risk among mammals: their phylogenetic proximity to *Homo sapiens*. We investigate how current extinction trends could modify our own position on the tree of life and the consequence for mammalian phylogenetic diversity and conservation if present-day trends continue. We then evaluate whether the observed nonrandom patterns of extinction risks could drive the future phylogenetic isolation (originality) of *H. sapiens*. A species is defined as original if it has few close relatives [10]. Currently, primates constitute one of the most species-rich orders of mammals. However, among primates, the family Hominidae contains seven species only, including *H. sapiens*, and all but one are threatened with extinction.

We also explore the reasons underlying the extinction risks of our close relatives, which can be placed within two broad categories: intrinsic factors linked to species biology and extrinsic factors representing the threats species are faced with in their environment. At the global scale, large home range, lower population growth [11], long generation times, low dispersal ability [12], high trophic level [13], habitat and diet specialism [14], and large body mass [11, 15, 16] have been identified as intrinsic factors associated with species sensitivity to threats in mammals [5]. Among these, large body mass has been considered as a potential catalyst of species sensitivity [11, 15]. The life-history traits that increase species’ vulnerability to anthropogenic threats frequently scale with body size.

Extrinsic factors may drive species extinctions independent of species traits or in combination with them. The International Union for Conservation of Nature (IUCN) Red List revealed that a high proportion of mammals, and of primates within mammals, are impacted by three types of threats: (1) exploitation, especially by hunting and trapping of terrestrial animals and indirectly by logging and wood harvesting; (2) agriculture; and (3) housing and urban areas [8, 9]. But whether threats are biased towards the species most closely related to *H. sapiens* has not been explored. Yet, species may be differently affected by threats not just because of their biology but also because of their ecological and cultural relationships with humans. For example, local human populations have developed different practices of hunting primates especially for food, traditional medicinal or cultural rituals [17-20]. In Africa, Eastern lowland Gorillas have been drastically affected by armed soldiers and war-ravaged refugees who make camps in the forest [20]. When hunting is accompanied by infant harvesting, apes (Hominidae, Hylobatidae) as pets have low rates of survivorship [21] and may thus be more affected. Also the accumulation of threats on a single species may explain its level of extinction risk [5]. We thus explored which and how many threats particularly affect the primates most closely related to *H. sapiens*.

In our analyses we ask the following questions: Are species that are most closely related to *H. sapiens* more at risk of extinction? Does mammal body mass, on average, tend to increase with phylogenetic proximity with *H. sapiens*? What are the major threats that act on and characterize the species most closely related to *H. sapiens*? Will *H. sapiens* become a sole survivor of its lineage and a phylogenetically isolated species?

## 2. Material and Methods

### (a) Mammalian extinction risk status

We have used the IUCN Red List [8] to classify each mammal species into one of the following categories: Least Concern (LC), Near Threatened (NT), Vulnerable (VU), Endangered (EN), Critically Endangered (CR). We excluded extinct species and those extinct in the wild. We considered species for which limited biological information was available or Data Deficient (DD) species separately. We obtained classifications for 5451 mammal species of which 435 were primates.

### (b) Threats

The IUCN Red List [8] contains a classification composed of 3 levels of direct threats [22]. The first and second levels are exclusive. Although all first-level threats are divided into second-level threats, they are not all divided into third-level threats. The third level is thus not necessarily complete as it contains detailed examples of threats [22]. The IUCN data base indicates whether the threats are ongoing, past or future. We considered ongoing threats in our analyses. 94% of threats affecting mammals were noted as ongoing with 97% ongoing in primates. 83% of primates had specified ongoing threats but only 50% of all mammals. Part of this difference could be due to the many poorly known mammal species. We thus removed species without identified threat when analyzing threat types.

### (c) Non-human mammal geographic ranges

Geographic distributions were obtained from the IUCN Red List [8]. Each primate species’ known range is available as a polygon or a set of polygons in case of fragmented distribution. We selected only polygons where the species is resident and is considered as extant, i.e. the species is known or thought very likely to occur presently in the area, which encompasses localities with current or recent (last 20-30 years) records with suitable habitat at appropriate altitudes remaining. The data are held in shapefiles, the ESRI native format. We used Mollweide projection to calculate the total area of these IUCN digital distribution maps for 5393 mammal species and 421 primate species.

### (d) Body mass

Body mass for each species was obtained from the PanTHERIA data base [23] complemented with the EltonTraits data base [24], which relies on Smith *et al.* [25]. We excluded interpolated data. We obtained body mass for 4479 species, among which 284 were primates.

### (e) Taxonomy and phylogeny

The taxonomy was available for all species in the IUCN Red List [8] following Wilson and Reeder [26]. For mammals, we used Bininda-Emonds *et al.* time tree [27] improved by Fritz *et al.* [15], referred to as Fritz *et al.* phylogeny. We also considered Rolland *et al.* [28] who resolved the Fritz *et al.* tree using the polytomy resolver of Kuhn *et al.* [29]. We refer to the resulting time tree as the Rolland *et al.* phylogeny. For primates, we pruned Fritz *et al.* and Rolland *et al.* trees conserving primate species only. We also considered the four molecular phylogenetic trees from Springer *et al.* [30]. Springer *et al.* time trees differed in their estimation of divergence times with the relaxed clock enforcing autocorrelated rates and hard-bounded constraints (AUTOhard), autocorrelated rates and soft-bounded constraints (AUTOsoft), independent rates and hard-bounded constraints (IRhard), or independent rates and soft-bounded constraints (IRsoft).

### (f) Clades defined by their distance to *H. sapiens*

The phylogenetic distance of a given species to *H. sapiens* was defined as the age of its first common ancestor with *H. sapiens*. For part of our analyses, we grouped species into monophyletic clades according to their distance to *H. sapiens*. Although the estimation of clade age may change, the delimitation of the clades was invariant across phylogenies.

### (g) Linking the phylogenetic distance to *H. sapiens* and species’ extinction risks

To evaluate whether species closely related to *H. sapiens* tend to be more threatened than expected randomly, we associated species on the mammalian [15] and primate phylogenetic trees [30] with the IUCN Red List data [8]. In each clade defined by its distance to *H. sapiens* (see previous section), we counted the number of species per extinction risk status: considering Least Concern, Near Threatened, Vulnerable, Endangered, Critically Endangered species, thus excluding Data Deficient species. We conducted a Monte Carlo χ^2^ test [31] with 10,000 simulations first for all mammals and then for primates.

### (h) Correlation between phylogenetic distance to *H. sapiens* and species’ body mass

The connection between the phylogenetic distance to *H. sapiens* and each non-human species body mass was analyzed with Spearman rank correlation. We considered only species for which we had information on both body mass and phylogeny (4395 mammals, 242 primates).

### (i) Threat analysis

We analyzed the ongoing threats identified in IUCN Red List classification [8]. We calculated: 1. the number of first-level threats, second-level threats and third-level threats per species in the mammalian clades defined by their distance to *H. sapiens*; 2. the mean and standard deviation of the phylogenetic distances between *H. sapiens* and the species affected by each second-level threat (the most detailed and complete level of threats). We used a Monte Carlo test to determine whether the average distance between *H. sapiens* and the species affected by a second-level threat was lower than expected by chance. For that we used the following steps: 1. we calculated the observed average distance between *H. sapiens* and the *N* species affected by a second-level threat (Obs.); 2. we randomly drawn *N* species among those with at least one specified threat (whatever the threat), and calculated the average distance (Sim.) between *H. sapiens* and these sampled *N* species; 3. we repeated step#2 1,000 times; 4. we calculated the P-value as the proportion of time Sim. was lower than or equal to Obs.

### (j) *H. sapiens*’ originality

We considered two indices of species’ phylogenetic originality: an index named “evolutionary distinctiveness” (*ED*) by Isaac *et al.* [32] and an index based on ‘quadratic entropy’, which we abbreviate as *Qb* [33]. *ED* distributes the phylogenetic diversity (sum of branch lengths) contained in a phylogenetic tree uniquely among the species at the tips. This is achieved by dividing the shared evolutionary history represented by a branch equally among its daughter species at the tips. The index *Qb* is the vector of species’ relative abundance that maximizes the quadratic entropy, an established measure of diversity defined in this context as the average phylogenetic distance between individuals from an assemblage of species. Compared to each other, *ED* is strongly dependent on the shortest phylogenetic distance between the focal species and all others while *Qb* is also sensible to the amount of past evolution shared by species and to the average phylogenetic distance between a species and all others in a set [5, 34].

For each phylogenetic tree (two for mammals and six for primates), and for each originality index, we ordered species from the most to the least original. In case of ties, we attributed the average rank. We performed this ranking by first retaining all species (no extinction) and then with successive scenarios of species extinction: we simulated the extinction of Critically Endangered species (Extinction scenario#1), followed by Endangered and Critically Endangered species (Extinction scenario#2), followed by Vulnerable, Endangered and Critically Endangered species (Extinction scenario#3), and then Near Threatened, Vulnerable, Endangered and Critically Endangered species combined (Extinction scenario#4). We evaluated the hypothesis that the originality of *H. sapiens* under each scenario of species extinction is higher than the originality expected if the extinction risks of mammals were random among the tips of the phylogenetic tree. For each scenario of species extinctions, we used a Monte Carlo approach to test this hypothesis, with the following steps: 1. we simulated species extinctions according to the selected scenario of extinction (for example, in Extinction scenario#1, all Critically Endangered species are driven extinct), and we calculated Obs. the originality rank for *H. sapiens* (ranking species from the most to the least original) among the surviving species; 2. we permuted the extinction risk status among all nonhuman mammals, simulated species extinctions according to the same selected scenario of extinctions and calculated the originality rank for *H. sapiens* (Sim.) among the surviving species; 3. we repeated step#2 500 times; 4. we calculated the P-value as the proportion of time Sim. was lower than or equal to Obs.

We performed the same analyses but on primates only using 1,000 permutations at step#3. As a first approach we excluded, from our analyses, species with missing phylogenetic positions. For Data Deficient species in the IUCN Red List, we considered extreme scenarios where these species were either all driven extinct or survived.

As a second approach, we randomly assigned phylogenetic positions to all phylogenetically unplaced species, and imputed extinction risk status to species classified as Data Deficient in the IUCN Red List. This was done to test the robustness of our results to missing data. In the phylogenetic trees, we randomly connected missing species to the smallest monophyletic clade (subtree) which contained all available species from their family. The probability that a new tip (here species) is added along any branch was directly proportional to the length of the branch. The order in which species were added was randomized. We repeated this operation 200 times for mammals and 500 times for primates. We ended up with the simulation of 200*2=400 complete trees for mammals, and 500*6=3000 complete trees for primates.

To impute extinction risk status for Data Deficient species, we collected for each species the extinction risk status and variables known to be strongly linked with probabilities of extinction: taxonomy (order when we considered all mammals, family when we focused on primates), body mass, and geographic range [11, 35, 36]. Then, we used the missForest algorithm to impute values [37]. The missForest algorithm was repeated 200 times for mammals and 500 times for primates to account for uncertainties in predictions.

We thus ended up with 400 data sets (200 repetitions of the randomly completed trees each associated with a completed vector of extinction risks for each core phylogenetic tree [Fritz et al. and Rolland et al. phylogenies]) with a phylogenetic tree and a vector of extinction risk status for 5451 mammal species, and 3000 data sets with a phylogenetic tree and a vector of extinction risk status for 435 primate species. These data sets are not estimations of true phylogenies and status but are tools to evaluate the robustness of our results to uncertainties due to missing data.

We applied the analysis above on *H. sapiens*’ originality to each data set reducing the number of permutations in Monte Carlo tests to 200 for mammals and 500 for primates given the high number of data sets treated and the large size of the mammal phylogeny. This led to the calculation of 320,000 vectors of originality for mammal species (400 data sets * 4 extinction scenarios * 200 permutations per scenario) and 6,000,000 vectors for primate species (3000 data sets * 4 extinction scenarios * 500 permutations per scenario).

All analyses were performed in R [38]. We used a nominal α level of 0.005 for all our tests.

## 3. Results

### (a) Extinction risk is bias towards species closely related to *H. sapiens*

We found higher differences in extinction risks between clades than expected by chance (χ^2^=692.8, P<<0.01). All primate clades tend to have more Endangered and/or Critically Endangered species than if extinction risks were randomly distributed among mammal species (table S1). Extinction risks among Dermoptera are close to random, while Scandentia, Rodentia and Lagomorpha have a higher proportion of Least Concern species than expected. The other large clades, more distant to *H. sapiens*, tend to have a slight excess of Near Threatened and/or Vulnerable species, compared to a random distribution of extinction risks. An exception is the endemic clade of Monotremata species, with five species only. It is the farthest clade to *H. sapiens*; it contains a significant proportion of Near Threatened species and Critically Endangered species. Within the primate phylogeny, we also found that clades with a recent common ancestor with *H. sapiens* were the most at risk (χ^2^=87.4, P<<0.01; figure 1; table S2). Our close relatives, the orangutans and gorillas (family Hominidae) are Critically Endangered, followed by, in order of threat, the common chimpanzee, the bonobo (family Hominidae), the gibbons (family Hylobatidae), the Strepsirrhini clade (lemuriform primates) and the tarsiers (Tarsiidae family). Conversely, species from the Cercopithecidae and the New World families are less threatened than expected. Overall, within mammals and within primates, the relationship between extinction risk and the distance to *H. sapiens* has a U-shape, with the clades most at risk being the closest to *H. sapiens*, the second most at risk the farthest, the least at risk having intermediate distances (figure 1; tables S1 and S2).

**Figure 1.**
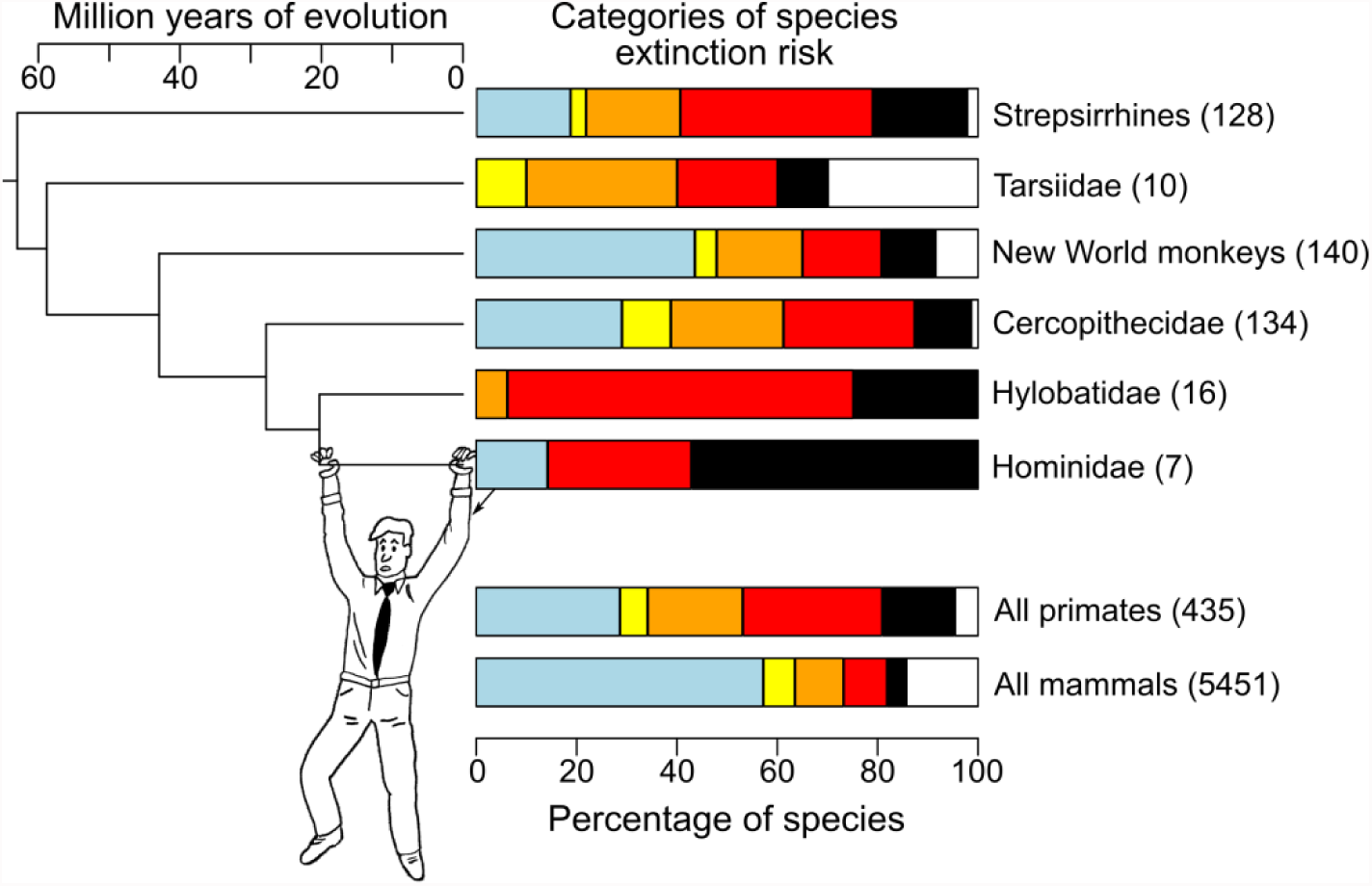
High extinction risks for primates, especially for families closely related to *Homo sapiens*. A partial representation of the primate phylogeny is given. It shows the relationships between *H. sapiens* (family: Hominidae) and other species grouped by monophyletic clades, according to their distance to *H. sapiens*, including the New World families (Aotidae, Atelidae, Callitrichidae, Cebidae, Pitheciidae) and Strepsirrhines (Cheirogaleidae, Galagidae, Indriidae, Lemuridae, Lepilemuridae, Lorisidae). Next to the phylogenetic tree, a bar plot gives the percentage of species defined as Least Concern (light blue), Near Threatened (yellow), Vulnerable (orange), Endangered (red), and Critically Endangered (black) [8]. The remaining percentage left in white represents species without an attributed category in the IUCN Red List (Data Deficient species). The number of species per clade evaluated in the IUCN Red List (including Data Deficient species) is given in brackets.

### (b) Intrinsic and extrinsic factors may explain the higher extinction risk of human close relatives

We found that species’ body mass decreases with the phylogenetic distance to *H. sapiens* (Spearman rank correlation r=-0.66, n=242, P<<0.01; figure S1) within primates, and to a lesser extent across all mammals (r=-0.25, n=4395, P<<0.01). We also found that species closely related to *H. sapiens*, especially the apes, tended to be affected by a higher number of threat types compared to other non-primate species (figure S2). Within primates, 11 out of 26 second-level threats impact species closely related to *H. sapiens* more frequently than expected by chance (figure 2). Among these 11 threats, some affect few species, with the exception of species closely related to *H. sapiens*, in particular viral and prion-induced diseases and habitat shifting and alteration due to climate change and severe weather. Others, like housing and urban areas and hunting affect many species, but more frequently species closely related to *H. sapiens* than expected by chance.

**Figure 2.**
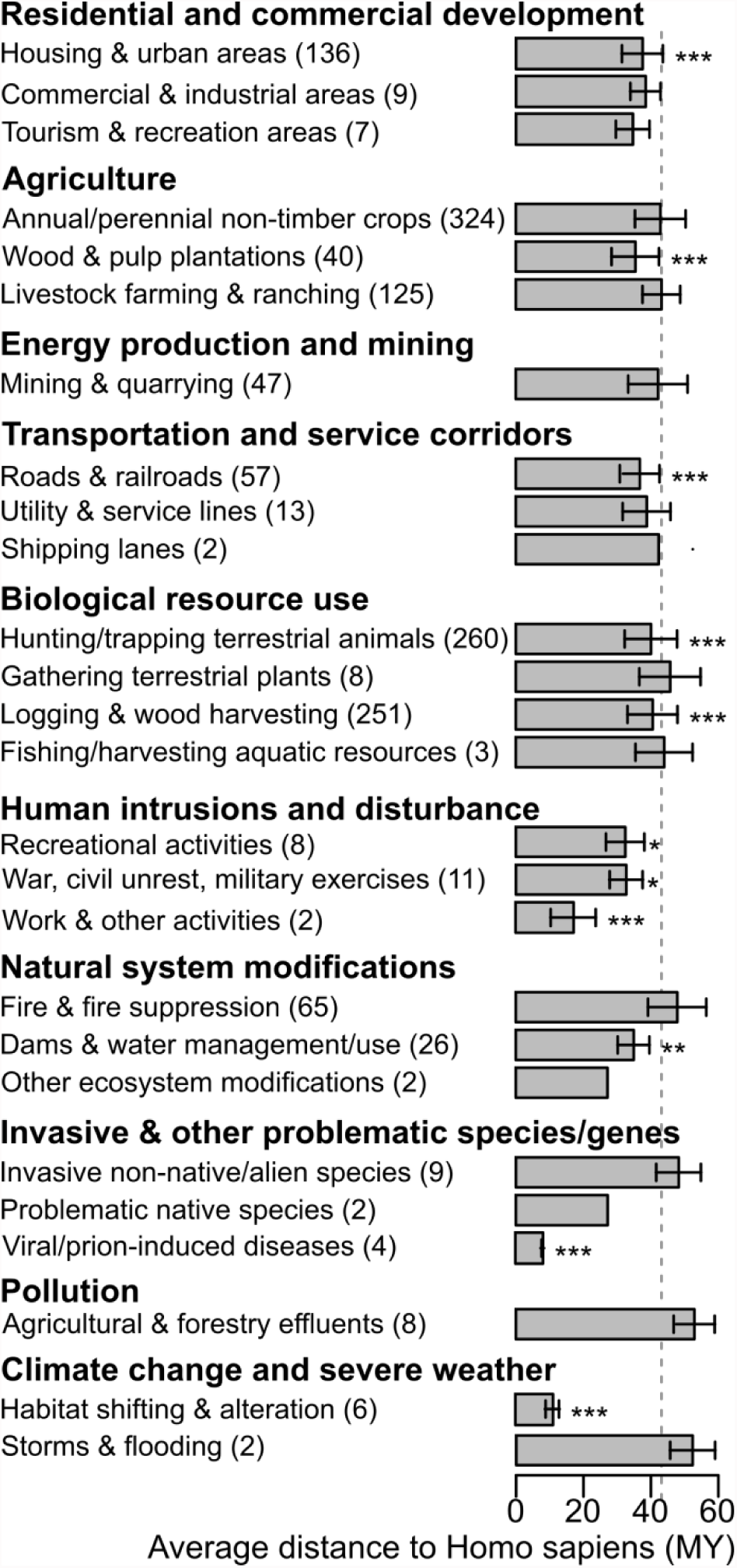
Link between threats and phylogenetic distance to *Homo sapiens* within primates. The number of species affected by each second-level threat is given in brackets. The bar plot provides the average phylogenetic distance between *H. sapiens* and primate species affected by each second-level threat. Only ongoing threats known to affect at least one primate species were included. Segments indicate standard deviation. The dashed grey line represents the average phylogenetic distance between a primate species with known ongoing threat(s) and *H. sapiens*. Monte Carlo test: *0.01<P≤0.05, **0.005<P≤0.01, ***P≤0.005.

### (c) *H. sapiens* is bound to become a phylogenetically isolated species

Results are summarized in table 1 for index *Qb*, Fritz *et al.* phylogeny for mammals and Springer *et al.* AUTOhard phylogeny for primates, and presented in detail, with all indices and phylogenies, in tables S3 to S6. We estimated the current originality rank for *H. sapiens* to be 460 among 5451 mammal species and 30 among 435 primate species. If all the threatened (Vulnerable, Endangered and Critically Endangered) species are lost, *H. sapiens* would rank among the 100 most original mammal species and among the five most original primate species (table 1). Such a dramatic alteration in rank would be highly unlikely if the threatened species were randomly distributed across the phylogeny (table 1).

**Table 1.** *Homo sapiens* originality rank. Results here are averaged over all our simulations, which account for missing data (detailed results in Tables S3 to S6). We first measured Obs., the observed rank for the originality of *H. sapiens*, ordering species from the highest to the lowest originality. Next we drove all Critically Endangered (CR) species to extinction, and performed the same calculation. We compared the observed rank for *H. sapiens* originality with ranks obtained permuting extinction risks between mammals (200 times) or primates (500 times). The average simulated rank (Sim.) for *H. sapiens* and its standard deviation (s.d.) are given. We repeated this approach driving Endangered (EN) and CR species, next Vulnerable (VU) to CR and then Near Threatened (NT) to CR species to extinction. Proportion of times Sim. was lower than or equal to Obs.: ***P ≤ 0.005.

| Extinctions | Nb. of species | Originality rank |  |
| --- | --- | --- | --- |
|  |  | Obs. | Sim. (s.d.) |
| Mammals |  |  |  |
| None | 5451 | 460 |  |
| CR | 5138 | 251*** | 440.40 (17.16) |
| EN, CR | 4456 | 90*** | 398.31 (35.92) |
| VU to CR | 3785 | 52*** | 352.73 (47.39) |
| NT to CR | 3373 | 44*** | 322.84 (50.45) |
| Primates |  |  |  |
| None | 435 | 30 |  |
| CR | 369 | 11*** | 27.59 (3.41) |
| EN, CR | 244 | 4*** | 21.17 (5.97) |
| VU to CR | 157 | 2*** | 16.29 (6.35) |
| NT to CR | 131 | 2*** | 14.70 (6.18) |

This result is robust to species for which there is inadequate information to make an assessment of their extinction risk and it is also robust to the estimation of the phylogenetic tree (tables S3 to S6). We chose to present results of *Qb* which synthesize more the whole shape of the phylogenetic tree when estimating the originality of a species. However, with *ED* the results are even stronger, reinforcing our conclusions: the predicted loss of all Hominidae species and almost all Hylobatidae species drastically increases *H. sapiens ED* value (tables S3 to S6).

According to our simulations, if all threatened primate species are lost, the only primate that would be always more phylogenetically original than *H. sapiens*, is the Near Threatened Phillipine tarsier (*Tarsius syrichta*) (table S7). If all threatened and near threatened primate species were driven extinct, only the currently Least Concern western fat-tailed dwarf lemur (*Cheirogaleus medius*) is ranked more phylogenetically original than *H. sapiens*, regardless of the originality index and the phylogeny used (table S7).

## 4. Discussion

Here, we have investigated the consequences of phylogenetic proximity to humans on primate extinction and human phylogenetic originality. Nonhuman primates have taken on particular places in human belief, mythology and cultural ecology [39], and this could have generated conservation conflicts. There is increasing evidence for a link between threat and phylogenetic proximity to humans affecting primate conservation. For instance, the transmission of disease has particular consequences as our closest primates can acquire our diseases but may not have the associated immune defenses, as unfortunately demonstrated within the great apes (Hominidae) [9]. However, there is no clear evidence that, in general, human activities deliberately target our closest relatives. The higher extinction risks of species closely related to *H. sapiens* thus likely stem from their exposure to a large variety of threat types and to their intrinsic greater sensitivity to threats.

Our results indeed show that body mass is inversely correlated with the phylogenetic distance to *H. sapiens* within mammals and within primates. In mammals, the correlation between extinction risk and body mass at small geographic scale was found to be significant only in the tropics [15]. Most primate species live in tropical forests [9]. Their range regions overlap with a large and rapidly growing population with high level of poverty [9]. Primates are affected by a diversity of threats resulting from local and global market demands, from hunting and illegal trades, from climate change and disease [9]. Body mass is correlated with a range of other traits such as large home range, specialism and lower population growth which might reduce ability to compensate for increased mortality. In particular our closest relatives, the great apes, have slow reproductive rate due to the long development time required for social learning (e.g. complex feeding behavior and social interactions, culture, environmental instability) [40, 41]. They may be more sensitive to the diversity of threats in their regions. The traits that helped make humans so successful may thus also be the ones that link to higher extinction risk, such as large body size, which may be associated with increasing brain size but also increasing development times and decreasing reproductive rates.

Given the high number of species at risk of extinction and limited conservation resources, we cannot avoid the ‘agony of choice’ and the opportunity costs of selecting which species or habitat to protect [42]. Among the many potential criteria used to prioritize conservation actions, the originality of a species is increasingly recognized as important as phylogenetically original species may represent rare biological features that few other species possess [32, 33, 43]. It is thus ironic that *H. sapiens* is likely to be a highly original species among living mammals in the near future, and one of the most original primate species. It is all the more ironic that our study shows we, humans, might be the driver of our own phylogenetic distinctiveness by endangering the existence and heightening the extinction risk of our closest mammalian relatives. Original species have often been likened to living fossils or taxonomic relicts. However, our study shows that high species originality can be a product of recent extinction, and thus the originality of a species is independent of its age [44]. *H. sapiens* are neither living fossils nor relicts. By pruning the tree of life around us, we are increasing our originality and thus increasing the rarity of our biological characteristics, as well as losing evolutionary insights into their origins.

Due to the high risk of extinction, designing appropriate conservation strategies for our close relatives is critically important, but requires consideration of the many costs to local peoples [39]. While primates act importantly as seed dispersers to regenerate and maintain the forest ecosystem [45], they can also act as disease reservoirs [46, 47] and as pests raiding crops [48] and even our homes [20]. The interactions between humans and non-human primates are numerous. As human cultures, needs and priorities constantly change, so do these interactions, challenging conservation actions that must constantly adapt to reflect these shifts. It is not immediately obvious, however, why we should care that we are becoming increasingly phylogenetically original; we are probably the mammal species least at risk of extinction.

The pruning of the primate tree started well before the emergence of *H. sapiens.* Apes were much more diverse in the Miocene than now. By the end of the Miocene, the Earth had become littered with extinct ape lineages [49]. The same trend is seen in hominins: *H. sapiens* is the sole member of a much larger evolutionary bush of hominin species [50]. Why other hominins went extinct is still controversial. One of the existing hypotheses is that *H. sapiens* has been already the cause of the extinction of other close relatives, such as *H. neanderthalensis* [51].

A recent study suggests that extinction rates throughout primate history tended to decline over time [52]. However, we suggest that the rhythm of primate extinctions is likely to change dramatically: our study shows accelerating primate extinctions with our closest relatives being the most likely to be lost first. Our close living relatives shed light on the origins of our diseases, on our own origins and adaptations, especially regarding our unique cognitive capacities, our behaviors (the most brilliant and the most peculiar ones, including spite and war [53]), our capacity of self-medication, to have culture, to communicate with complex language, and much more that we cannot discern from fossils. Their loss is our loss.

Through our actions we are isolating ourselves. Our study shows the future evolutionary trajectory of our loneliness. With the loss of our close relatives, we lose not only unique biodiversity that is essential to maintain ecosystem functions and the Earth climate but also a sense of our own fragility, our connection with the environment, reinforcing our delusions of success. Our actions blur the question of what makes us distinct, what makes us “humans”. Extinctions will leave us without a mirror to contextualize our biology. They accentuate the misconception that we are unique and then, ironically lead us to fulfill it.

## Data

We accessed and downloaded data from the IUCN Red List on 04 October 2016 for the extinction risk categories, on 05 October 2016 for digital distribution maps and on 22 January 2017 for threats, http://www.iucnredlist.org/.

## Acknowledgments

We thank Giuseppe Donati, Rob Dunn, Dieter Lukas, Arne Mooers and Carlo Ricotta for their comments on our paper.

